# Development and optimisation of a high-throughput screening assay for in vitro anti–SARS-CoV-2 activity: evaluation of 5676 phase 1 passed structures

**DOI:** 10.1101/2022.02.02.478671

**Authors:** Winston Chiu, Lore Verschueren, Christel Van den Eynde, Christophe Buyck, Sandra De Meyer, Dirk Jochmans, Denisa Bojkova, Sandra Ciesek, Jindrich Cinatl, Steven De Jonghe, Pieter Leyssen, Johan Neyts, Marnix Van Loock, Ellen Van Damme

## Abstract

Although vaccines are currently used to control the coronavirus disease 2019 (COVID-19) pandemic, treatment options are urgently needed for those who cannot be vaccinated and for future outbreaks involving new severe acute respiratory syndrome coronavirus virus 2 (SARS-CoV-2) strains or coronaviruses not covered by current vaccines. Thus far, few existing antivirals are known to be effective against SARS-CoV-2 and clinically successful against COVID-19.

As part of an immediate response to the COVID-19 pandemic, a high-throughput, high content imaging–based SARS-CoV-2 infection assay was developed in VeroE6-eGFP cells and was used to screen a library of 5676 compounds that passed phase 1 clinical trials. Eight candidates (nelfinavir, RG-12915, itraconazole, chloroquine, hydroxychloroquine, sematilide, remdesivir, and doxorubicin) with *in vitro* anti–SARS-CoV-2 activity in VeroE6-eGFP and/or Caco-2 cell lines were identified. However, apart from remdesivir, toxicity and pharmacokinetic data did not support further clinical development of these compounds for COVID-19 treatment.

## INTRODUCTION

Coronaviruses (CoVs) are enveloped, positive-sense single-stranded RNA viruses; of the 7 members of the CoV family known to infect humans, most cause mild respiratory disease.^1^ In the last 2 decades, betacoronaviruses have caused outbreaks of severe respiratory disease, including severe acute respiratory syndrome (SARS) in 2002 and 2003, caused by SARS-CoV-1,^2^ followed by Middle Eastern Respiratory Syndrome (MERS) in 2012, caused by MERS-CoV.^3^ In late 2019, an outbreak of a novel respiratory syndrome, coronavirus disease 2019 (COVID-19), was reported in Wuhan, China.^4^ Common presenting symptoms of COVID-19 caused by Alpha variants include cough, fever, loss of taste or smell, and fatigue.^5^ However, the more recent Delta variant seems to present slightly different symptoms, such as headache, runny nose, throat ache, fever, and coughing.^5^ In cases that progress to severe disease, patients commonly experience dyspnoea and hypoxaemia followed by respiratory failure.^6^ COVID-19 and its etiologic agent, severe acute respiratory syndrome coronavirus 2 (SARS-CoV-2), have spread globally since the initial outbreak, infecting >210 million people and leading to ≥4 million deaths.

Even before the World Health Organization declared COVID-19 a public health emergency of international concern on 30 January 2020, repurposing of existing drugs and drug candidates was explored to accelerate the traditional research and development timelines in order to provide a rapid response to this unmet medical need.^7^ Drug repurposing had been applied previously for SARS-CoV-1, MERS-CoV, and other viruses.^7,8^ Early in the pandemic, development of cell-based systems was critical for the rapid evaluation of drugs with antiviral activity against SARS-CoV-2. Similar to SARS-CoV-1, Vero cells and Caco-2 cells were found to be susceptible to infection with SARS-CoV-2.^9,10^ Therefore, a previously published high-throughput screening (HTS) assay for SARS-CoV-1^11^ using authentic infection of VeroE6 African green monkey kidney epithelial cells expressing a stable enhanced green fluorescent protein (VeroE6-eGFP) was adapted, further developed, and miniaturised for screening of antiviral drugs against SARS-CoV-2, isolated from a Belgian patient. The resulting assay is an HTS 384-well cell-based SARS-CoV-2 infection assay with a high content imaging (HCI) readout of fluorescence that provides a measure for cytopathic effect (CPE). In parallel, a cellular toxicity assay was developed using ATPlite™; any toxicity of the compound to the cell line is then evaluated by luminescence.

To identify potential candidates for rapid clinical development, 5676 chemical structures that had passed phase 1 clinical studies with applications in a variety of therapeutic fields including oncology, neuroscience, and infectious diseases, were screened. The compounds were evaluated in a 7-point dose-response curve, starting at 20 µM, for their ability to inhibit SARS-CoV-2 induced CPE without causing general cellular toxicity. Selected hits were further evaluated in a second cellular model using Caco-2 cells that are susceptible to SARS-CoV-2 infection.^12,13^

## MATERIALS AND METHODS

### Assembly of the library

Janssen Pharmaceutica maintains a regularly updated database of compounds that have been approved or tested in a successfully completed clinical phase 1 study. One of the many uses of this **P**hase **O**ne **P**assed **S**tructures (**POPS**) database is to serve as a starting point to identify high-priority compounds for possible repurposing. The goal of the POPS database is to be highly enriched in druggable, well-documented, and diverse compounds from all disease areas, including oncology, neuroscience and infectious diseases. The database is updated on a regular basis, both from a (virtual) annotation standpoint and with a physically available set of compounds that can be screened. Due to previous acquisition and synthesis efforts and active purchasing in the first months of 2020, approximately 5500 compounds that had sufficient availability and passed quality control for purity were identified and plated.

### Plate production

Plates were freshly prepared to ensure high-quality assays and were submitted for screening. CELLSTAR^®^ 384-well plates (Greiner Bio-One, Vilvoorde, Belgium) were prespotted with a 300 nL compound in dimethyl sulfoxide (DMSO; 300 nL 100% DMSO for control wells) and each plate was spotted in triplicate. An Echo^®^ 555 Liquid Handler (Labcyte Inc., Indianapolis, USA) was used to spot the compounds at a final concentration of 20 µM. Spotted plates were frozen and transported to the Rega Institute of the KU Leuven, Belgium.

### Cell cultures

VeroE6-eGFP were cloned and validated in-house as previously described.^11^ Cells were cultured and maintained in Dulbecco’s modified Eagle medium (DMEM; Gibco, Waltham, USA) supplemented with 10% (vol/vol) heat-inactivated foetal bovine serum (FBS; Biowest, Riverside, USA), 0.75% sodium bicarbonate (Gibco), and 50 U/mL penicillin-streptomycin (Gibco). Caco-2 cells (human colon carcinoma cell line; obtained from the Deutsche Sammlung von Mikroorganismen und Zellkulturen, Braunschweig, Germany) were cultured in a minimal essential medium supplemented with 10% FBS with penicillin (100 IU/mL) and streptomycin (100 μg/mL). Cells were maintained at 37°C in 5% CO_2_.

### SARS-CoV-2 preparation

SARS-CoV-2-Belgium (strain BetaCov/Belgium/GHB-03021/2020) was recovered from a nasopharyngeal swab taken from an asymptomatic patient returning from Wuhan, China. Virus stocks were inoculated and passaged first in HuH-7 cells and then 5 times in VeroE6-eGFP cells prior to storage at –80°C. Passage 6 was used for all VeroE6-eGFP experiments (viral titre: 3.0 × 10^6^ TCID_50_/mL). All manipulations were performed in a licenced and certified biosafety level 3 (BSL-3) facility at the Laboratory of Virology & Chemotherapy at the Rega Institute of the KU Leuven, Belgium.

SARS-CoV-2/FFM1 (strain hCoV-19/Germany/FrankfurtFFM1/2020) was isolated from a German patient sample and passaged twice in Caco-2 cells prior to storage at –80°C. All passages were sequenced using a MinION platform (Oxford Nanopore). Viral titres were determined for the last 3 passages by end-point dilution assays. All manipulations were performed in a licenced and certified biosafety level 3 (BSL-3) facility at the Laboratory Medizinische Mikrobiologie at the Universitats Klinikum Frankfurt (Goethe University) in Germany.

### Antiviral and toxicity assays in VeroE6-eGFP cells

Each compound from the library was evaluated for antiviral activity using a cell-based VeroE6-eGFP assay. Two assays were developed simultaneously: the first one used whole well fluorescence measured with a multimode plate reader (PR; Tecan, Infinite M1000 Pro) and the other used HCI (Thermofisher, Arrayscan XTI). Spotted 384-well plates were seeded with 30 µL with 2000 (PR assay) or 8000 (HCI assay) VeroE6-eGFP cells in DMEM 2% FBS in each well.

An additional 30 µL medium was added to the cell controls. The plates used in the HCI assay were incubated overnight or approximately 20 hours prior to infection in a humidified incubator at 37°C with 5% CO_2_. After cell seeding (PR assay) or overnight incubation (HCI assay), the plates were transferred to the BSL-3+ Caps-It robotics system where 30 µL SARS-CoV-2 was added using a noncontact liquid handling system (Tecan, EVO 100) to achieve a multiplicity of infection (MOI) of 0.001. After infection, the plates were automatically transferred to the system’s integrated incubators (37°C with 5% CO_2_) for 5 (PR assay) or 4 (HCI assay) days before performing the readout (**Figure 1**).

**Figure 1.**
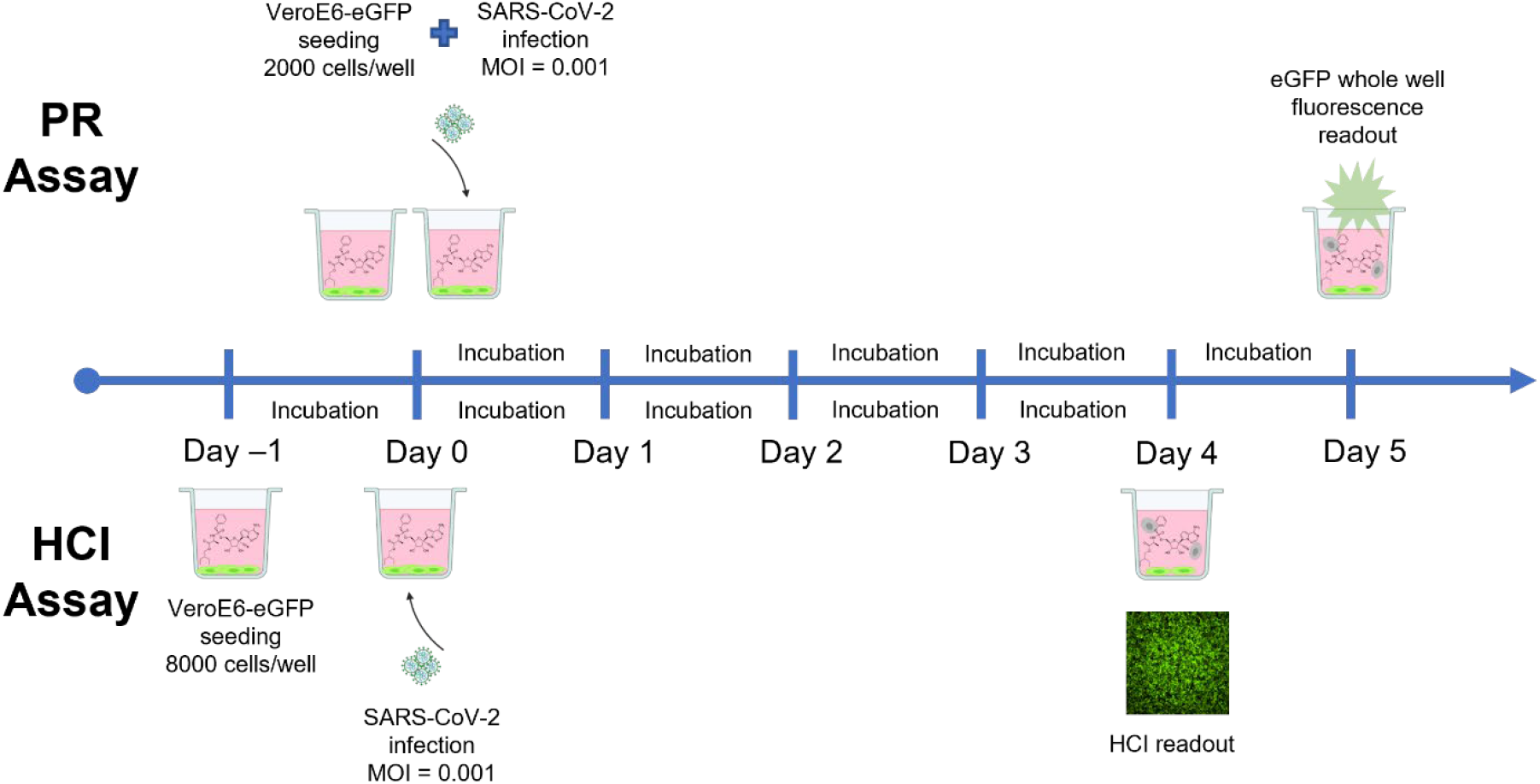
Schematic overview of PR and HCI assays using prespotted 384-well plates with compounds in a dose-response curve format at a final starting concentration of 20 µM. PR assay (top): 2000 cells/well were seeded on Day 0 and infected on the same day, after which a PR readout was performed on Day 5. HCI assay (bottom): 8000 cells/well were seeded on Day –1 and infection with SARS-CoV-2 was performed on Day 0, after which an HCI readout was performed on Day 4. Plates were incubated at 37°C and 5% CO_2_ between days. eGFP, enhanced green fluorescent protein; HCI, high content imaging; MOI, multiplicity of infection; PR, plate reader; SARS-CoV-2, severe acute respiratory syndrome coronavirus 2; VeroE6-eGFP, VeroE6 African green monkey kidney epithelial cells expressing a stable enhanced green fluorescent protein.

Whole well fluorescence settings on the plate reader were set on 488 nm excitation and 507 nm emission bandwidths with 20 µs integration time and 4 reads per well. PR data were obtained in .csv format. Image acquisition on the high-content imagers was set to 485/20 nm excitation wavelength with an exposure time of 23 ms. The emission signal was captured with a multiband BGRFRN filter set and a dichroic mirror using widefield microscopy technology. One field per well was imaged using a 5× objective with 2 × 2 binning and 1104 × 1104 pixel resolution. An image analysis was performed on the acquired images using the HCS studio software. A custom image analysis protocol was developed based on the Spot Detector BioApplication. A background correction was performed to remove nonspecific signal from the raw image files. With the entire well selected as the region of interest, a fixed threshold for pixel intensity was set to measure eGFP signal. SARS-CoV-2 induced CPE in VeroE6-eGFP cells, leading to a marked reduction of eGFP signal. In contrast, a strong eGFP signal was observed under conditions without the virus. A quality control assessment and data analysis were performed using several output features: SpotTotalAreaCh2, SpotTotalIntensityCh2, and Valid Object Count. Following assay completion, plates were automatically decontaminated by the Caps-It system.

Compounds were screened in dose-response, and dose-response curves plotting normalised values versus compound concentrations were generated in Genedata Screener^®^ (version 17.0.6 – Standard). A smart fit strategy with automatic model selection without automatic masking was used for plotting the graphs and calculating EC_50_ (concentration of the compound that inhibited 50% of the virus-induced CPE) values. A low threshold for hit selection was determined to minimise the risks of having false negatives. Only the whole well fluorescence data and the HCI output feature “SpotTotalAreaCh2” were used for calculation. When 1 of 3 replicate data points deviated, it was masked and not taken into consideration for calculations or fitting.

The PR assay was developed first, and data analysis could be performed on the extracted data using the already established informatics tools of Genedata Screener^®^. The HCI assay was developed during the initial screening using the PR assay. Since the extracted HCI data were more robust and accurate, the HCI assay was used instead of the PR assay during this screening campaign. Due to the urgency of the screen, data points of >900 approved (based on the robust Z’ factor [RZ’]) assay plates were already collected using the PR assay.

Toxicity was assessed using the ATPlite™ kit by measuring luminescence after a reaction of ATP with added D-luciferin and luciferase. 40 µL of VeroE6-eGFP cells were seeded at 2000 cells/well in pre-spotted 384-well plates. The plates were then placed for 5 days in a humidified incubator with 5% CO_2_. After incubation, steps were followed according to the ATPlite™ manufacturer’s instructions. The luminescence read-out was performed on the ViewLux™ plate reader (PerkinElmer). Data were analysed by non-linear curve fitting (four parameter fit) from a dose-response curve using GraphPad Prism to calculate CC_50_ (cytotoxic concentration of the compound that reduced cell viability to 50%).

### Antiviral and toxicity assay in Caco-2 cells

After compounds with potential antiviral activity in VeroE6 cells were identified, confirmation of their inhibition of virus-induced CPE was performed in Caco-2 cells. Caco-2 cells were cultured for 72 hours on 96-well plates (50,000 cells/well) and infected with SARS-CoV-2/FFM1 at an MOI of 0.01. After a 48-hour incubation of the virus, cells, and compound in MEM supplemented with 1% FBS, the CPE was visually scored by 2 independent laboratory technicians. Optical densities were measured at 560/620 nm in a Multiskan Reader. Cell viability in Caco-2 cells in the absence of virus was assessed using the 3-(4,5-dimethylthiazol-2-yl)-2,5-diphenyltetrazolium bromide (MTT) assay, as previously described.^14^ Data were analysed by 4-parameter curve fitting from a dose-response curve using GraphPad Prism to calculate the EC_50_ and CC_50_ (cytotoxic concentration of the compound that reduced cell viability to 50%).

### Development and optimisation of a high-throughput screening pipeline for both assays

Assay quality performance (inter- and intra-plate data comparison) and optimisation (cell density, viral input, incubation time, Z-prime, signal/noise) were performed on 384-well plates spotted with reference compounds. Raw data was normalised using the following formula and was used for most metrics:

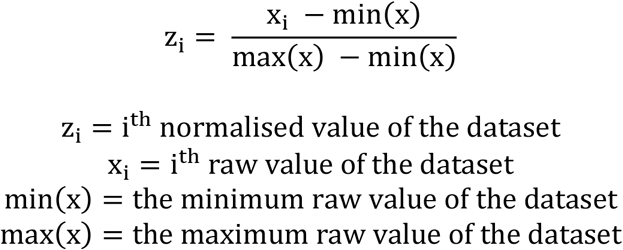

The Z-prime value, which encompasses the dynamic range of the assay and well-to-well variability, was calculated for each plate as a measure of assay quality.^15^ All compounds were screened in a dose response (7 dilutions) starting from 20 µM with half dilution steps and tested in triplicate on separate plates.

## RESULTS

### High-throughput assay development

At the start of the pandemic, an immediate public health response was needed but tools to screen for antiviral agents were not available at that time. Because of the high sequence homology between SARS-CoV-1 and SARS-CoV-2, the primary focus was on VeroE6 cells, which were previously shown to be readily susceptible to infection with SARS-CoV-2.^9,10^ Indeed, SARS-CoV-2 infected the cells and caused pronounced CPE, especially once the virus had been adapted to cell culture (3-5 passages on VeroE6 cells). Next, a SARS-CoV-1 antiviral assay in VeroE6-eGFP cells^11^ was adapted to be used for screening with SARS-CoV-2 (**Figure 1**). In this system, SARS-CoV-2 infection causes cellular CPE and loss of fluorescent signal. Compound-mediated antiviral activity reduces CPE, leading to increased fluorescence compared with controls. To monitor CPE and inhibition thereof by antiviral molecules, the assay was developed in parallel tracks whereby readout was performed either by fluorescence (using a GFP PR) or HCI at low resolution to visualise and quantify CPE (**Figure 1**).

In uninfected (cell control, CC) conditions, the mean normalised signal for the PR readout was 79.8 ± 10.9 (%CV = 13.7); for the HCI readout, the mean normalised signal was 94.8 ± 2.41 (%CV = 2.55). Under infected (virus control, VC) conditions, a high %CV was calculated for both readouts with a large variation between the data points; because the mean value was low (PR < 5 and HCI < 1), the %CV was highly sensitive to small changes. In this type of biological assay, unavoidable “eGFP cell debris,” found in residual organic waste after virus-induced cell death, contributes to variation in eGFP. For HTS quality assessment, the RZ’ and signal to noise ratio (S/N) were calculated for each method (PR readout: mean RZ’ 0.51 and S/N 27.3; HCI readout: mean RZ’ 0.91 and S/N 362) (**Figure 2**).

**Figure 2.**
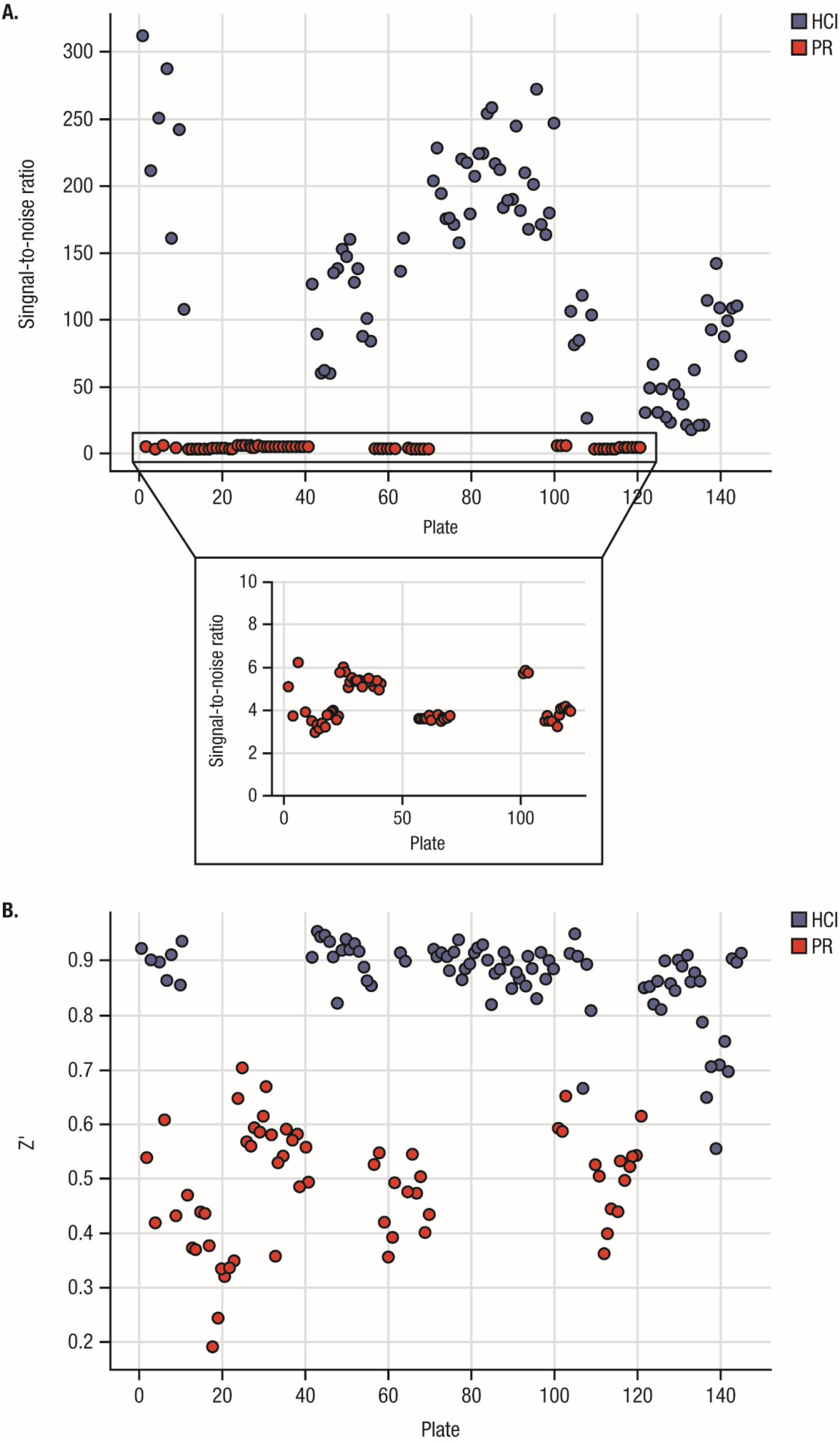
Signal-to-noise (**A**) and Z’ (**B**) comparison between PR (red) and HCI (blue) assays. The signal-to-noise ratio (raw data CC/raw data VC) for the PR assay ranged from 1.84 to 6.22 with Z’ values ranging from –0.26 to 0.72. The HCI assay yielded much higher values for both parameters; the signal-to-noise ratio ranged from 17.95 to 312.3 while Z’ values ranged from 0.57 to 0.95. CC, uninfected conditions; HCI, high content imaging; PR, plate reader; VC, infected conditions.

Reference compounds were used to verify the inhibition of SARS-CoV-2 infected in VeroE6-eGFP cells. Twelve compounds were selected based on what was known at the time regarding activity against SARS-CoV-1, MERS, and SARS-CoV-2: chloroquine,^8,16–18^ cinchocaine,^19^ colchicine,^20^ hydroxychloroquine,^21^ ponatinib,^22^ indomethacin,^22^ loperamide,^8^ lopinavir,^8^ nelfinavir,^23^ posaconazole,^24^ remdesivir,^17,18^ and saperconazole.^24^ These compounds were spotted 200 times more concentrated in quadruplicate at 300 nL/well, with a final concentration per well ranging from 10 µM to 100 µM depending on the compound, followed by a 7-point dilution with a dilution factor of 2. The EC_50_ value was calculated based on the obtained values from the HCI and PR readouts for each compound using the GeneData screener as described in the Methods.

Intra- and inter-plate comparisons were evaluated under uninfected conditions (CC) and SARS-CoV-2 infected conditions (VC) for both PR and HCI readout methods. In this step, plates containing reference compounds were seeded with VeroE6-eGFP cells as previously described for the PR and HCI antiviral assays to obtain CPE-induced eGFP signal reduction after SARS-CoV-2 infection. Intra-plate variation was evaluated by calculating the mean eGFP output signal and standard deviation (SD); these values were calculated from one plate under 32 uninfected conditions and 16 infected conditions. Inter-plate variation values were calculated using the data from all 4 individual experiments (4 plates; CC, 128 data points; VC, 64 data points). To assess reproducibility, measurements were obtained in 4 independent experiments. Additionally, the RZ’ and S/N were calculated for each experiment to determine HTS quality. Assay performance of both the PR and HCI antiviral assays are shown in **Table I**.

**Table I.** Assay performance of PR and HCI methods

|  |  | Intra-plate repeatability |  |  |  |  |  |  |  | Inter-plate reproducibility |  |
| --- | --- | --- | --- | --- | --- | --- | --- | --- | --- | --- | --- |
|  |  | Experiment 1 |  | Experiment 2 |  | Experiment 3 |  | Experiment 4 |  |  |  |
|  |  | PR | HCI | PR | HCI | PR | HCI | PR | HCI | PR | HCI |
| N |  | 32 | 32 | 32 | 32 | 32 | 32 | 32 | 32 | 128 | 128 |
| Uninfected cells (CC) | **Mean** | 79.2 | 93.3 | 81.1 | 93.6 | 82.4 | 96.6 | 76.3 | 95.6 | 79.8 | 94.8 |
|  | **SD** | 11.5 | 2.46 | 9.56 | 2.79 | 9.49 | 1.71 | 13.0 | 2.69 | 11.1 | 2.79 |
| N |  | 16 | 16 | 16 | 16 | 16 | 16 | 16 | 16 | 64 | 64 |
| SARS-CoV-2 infected cells (VC) | **Mean** | 2.86 | 0.16 | 2.24 | 0.26 | 4.30 | 0.37 | 2.95 | 0.40 | 3.09 | 0.30 |
|  | **SD** | 1.88 | 0.11 | 1.08 | 0.26 | 3.22 | 0.16 | 1.58 | 0.18 | 2.18 | 0.20 |
| Plate metrics | **RZ'** | 0.50 | 0.92 | 0.59 | 0.90 | 0.54 | 0.93 | 0.42 | 0.90 | 0.49 | 0.90 |
|  | **S/N** | 5.05 | 584 | 7.84 | 364 | 5.08 | 258 | 3.69 | 240 | 5.42 | 319 |

CC, cell control; HCI, high content imaging; PR, plate reader; RZ', robust Z' factor; SARS-CoV-2, severe acute respiratory syndrome coronavirus 2; SD, standard deviation; S/N, signal to background; VC, virus control.
Metrics were performed on normalised data, representing the percentage of green fluorescent signal.

Among the 12 compounds selected for assay development, 6 resulted in a dose-response relation on both readouts: chloroquine, cinchocaine, hydroxychloroquine, lopinavir, nelfinavir, and remdesivir. The mean and SD of the computed EC_50_ values from 16 replicates (4 intra-plate replicates and 4 plate replicates) and the toxicity data (CC_50_) computed from 5 replicates are shown in **Table II**. Cinchocaine (PR: 59.6 µM vs HCI: 63.2 µM), hydroxychloroquine (PR: 13.5 µM vs HCI: 17.7 µM), nelfinavir (PR: 3.5 µM vs HCI: 4.2 µM) and remdesivir (PR: 3.6 µM vs HCI: 2.6 µM) showed comparable EC_50_ values with no significant differences between both readouts (*P*>0.05). Chloroquine and lopinavir showed higher discrepancies between PR and HCI readouts, which can likely be attributed to physical differences (whole well fluorescence vs image analysis) and experimental differences (PR: 2000 cells/well, seeding and infection on day 0, reading 5 dpi; HCI: 8000 cells/well, seeding on day 1 and infection on day 0, reading 4 dpi). Remdesivir exhibited *in vitro* inhibition of SARS-CoV-2 with minimal effect on cell viability and was further used as a positive control.

**Table II.**
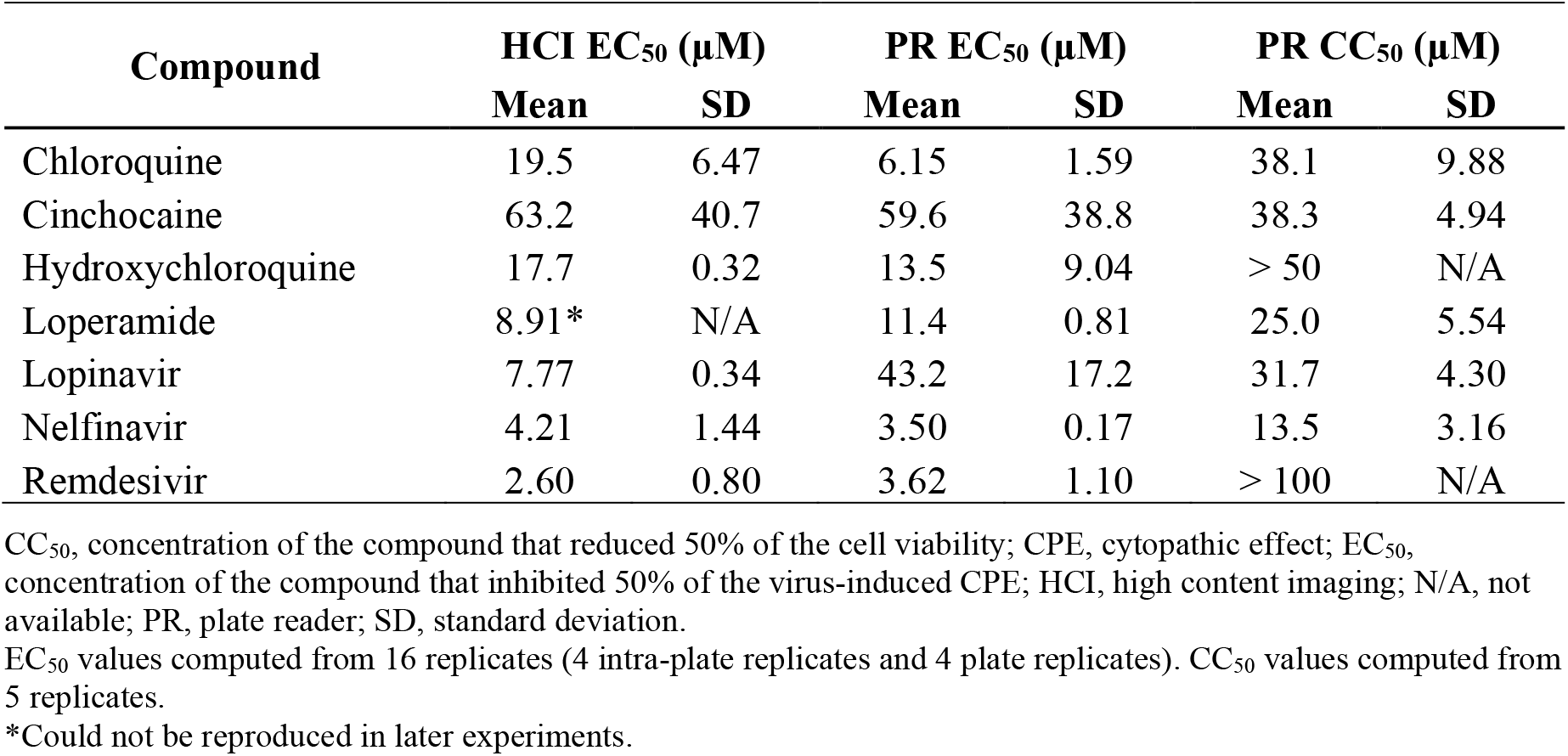
Mean antiviral activity (EC_50_) and cytotoxicity (CC_50_) values of reference compounds used for assay development

### Drug repurposing screen

Slightly cytotoxic compounds may result in false negatives in this assay (ie, activity caused by interference with cell proliferation). Therefore, all compounds were evaluated using dose-response curves, in which bell-shaped curves indicate upcoming cytotoxicity. In addition, every compound was tested in a parallel toxicity assay.

The drug repurposing screening hit rate was approximately 4.0%. All compounds were screened at the highest concentration available in 100% DMSO stock, which was either 100µM, 50µM or 25µM final concentration. Of the 5676 compounds screened, 228 compounds were found to be active (EC_50_ value <20 µM in either PR or HCI readout) (**Figure 3** and **Supplementary Table I**). All compounds were redissolved from neat and were tested again in 7-point with ½ dilution steps dose-response (starting at the highest possible concentration: 100µM, 50µM, or 25µM, final concentration) in triplicate for confirmation. In the confirmation run, 52 compounds remained active (maximum percent inhibition ≥30%). A thorough evaluation of these hits was made based on quality of the curve, the chemistry, potential/known mode of action, literature, and toxicity to select only those with potential clinical relevance.

**Figure 3.**
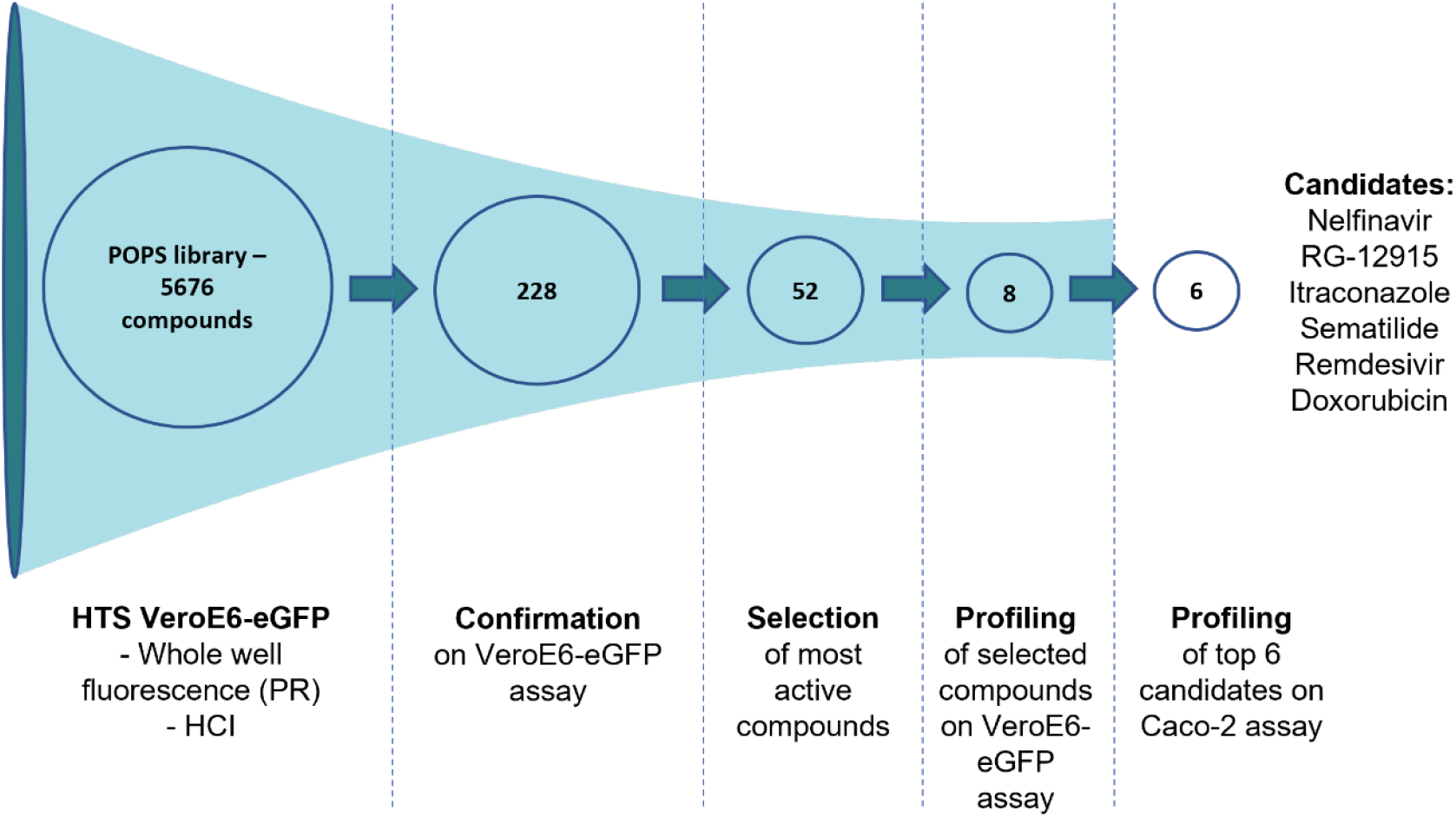
Schematic overview of the screening campaign used on the Janssen POPS library containing 5676 compounds. The high-throughput screen was performed on 384-well plates with the described PR or HCI VeroE6-eGFP assay. After the initial screening, 228 compounds showed activity against SARS-CoV-2 with EC_50_ values <20 µM in either assay. These compounds were retested in triplicate in the same assay for confirmation, which yielded 52 active compounds. Finally, the 8 most promising compounds were selected for further profiling, and the top 6 candidates were selected for additional testing using the Caco-2 assay. EC_50_, concentration of the compound that inhibited 50% of the infection; HCI, high content imaging; POPS, Phase One Passed Structures; PR, plate reader; VeroE6-eGFP, VeroE6 African green monkey kidney epithelial cells expressing a stable enhanced green fluorescent protein.

Finally, 8 compounds with antiviral activity against SARS-CoV-2 in the VeroE6-eGFP cells were selected for further investigation: chloroquine, doxorubicin, hydroxychloroquine, itraconazole, nelfinavir, remdesivir, RG-12915, and sematilide (**Table III**).

**Table III.** Observed antiviral activity (EC_50_) against SARS-CoV-2 and cytotoxicity (CC_50_) of the 8 compounds identified using the VeroE6-eGFP cell assay^a^

| **Compound** | **Class** | **EC<sub>50</sub>, μM**<br>**(min, max)** | **Repeats** | **CC<sub>50</sub>, μM**<br>**(min, max)** | **Repeats** |
| --- | --- | --- | --- | --- | --- |
| Sematilide | Antiarrhythmic | 3.9 (N/A) | 1 | >20 (N/A) | 1 |
| Remdesivir | Nucleotide analogue | 5.4 (1.15, 19.7) | 18 | >100 (>100, >100) | 10 |
| Nelfinavir | Protease inhibitor | 5.7 (3.29, >50) | 18 | 12.9 (10.3, 13.7) | 10 |
| Hydroxychloroquine | Antimalarial | 6.7 (N/A) | 1 | >50 (N/A) | 1 |
| RG-12915 | 5-HT<sub>3</sub> receptor antagonist | 6.8 (N/A) | 1 | 19.7 (N/A) | 1 |
| Chloroquine | Antimalarial | 7.3 (3.13, >50) | 12 | 35.1 (29.9, 50.3) | 10 |
| Doxorubicin | Anthracycline | 8.4 (5.93, 10.9) | 4 | >20 (N/A) | 1 |
| Itraconazole | Antifungal | >100 (5.13, >100) | 8 | >100 (>100, >100) | 8 |

5-HT<sub>3</sub>, 5-hydroxytryptamine; CC<sub>50</sub>, cytotoxic concentration of the compound that reduced cell viability to 50%; CPE, cytopathic effect; EC<sub>50</sub>, concentration of the compound that inhibited 50% of the virus-induced CPE; HCI, high content imaging; N/A, not available; SARS-CoV-2, severe acute respiratory syndrome coronavirus 2.
<sup>a</sup>Activity readout of the assay was based on HCI of fluorescent virus; cell toxicity was measured using ATPlite in uninfected cells.

### Confirmation of the top 6 hits in a Caco-2 infection assay

To exclude cell line–specific modes of antiviral activity, 6 compounds were further evaluated using a Caco-2 infection assay. Hydroxychloroquine and chloroquine were excluded from this evaluation because they had previously been found inactive in SARS-CoV-2–infected Caco-2 cells.^25^ This system has been used previously for SARS-CoV-1^11^ and was recently deployed for another drug repurposing screen.^25^ All compounds were evaluated with a dose-response curve to obtain an EC_50_ value based on visual inspection of CPE. In parallel, a CC_50_ value was obtained using a similar assay but without viral infection. Remdesivir and itraconazole exhibited a higher potency in Caco-2 cells compared to Vero cells; other compounds showed similar EC_50_ values in both cell lines. Sematilide was not active in Caco-2 cells. Dose-response curves of CPE and cytotoxicity readouts are shown in **Figure 4**; EC_50_ and CC_50_ values are summarised in **Table IV**.

**Figure 4.**
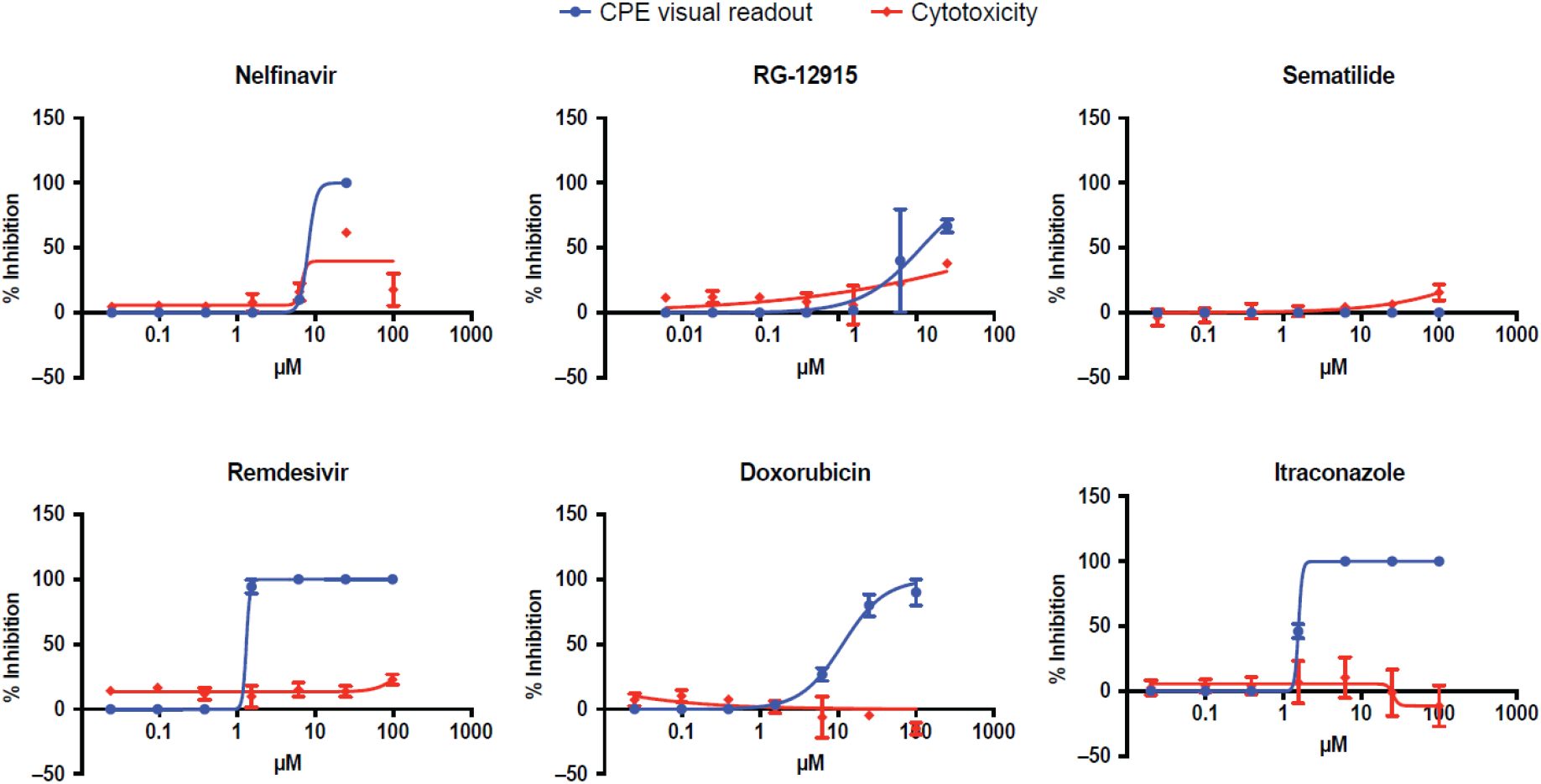
Effect on CPE assays and viability of Caco-2 cells of identified compounds. CC_50_, cytotoxic concentration of the compound that reduced cell viability to 50%; CPE, cytopathic effect; EC_50_, concentration of the compound that inhibited 50% of the virus-induced CPE.

**Table IV.**
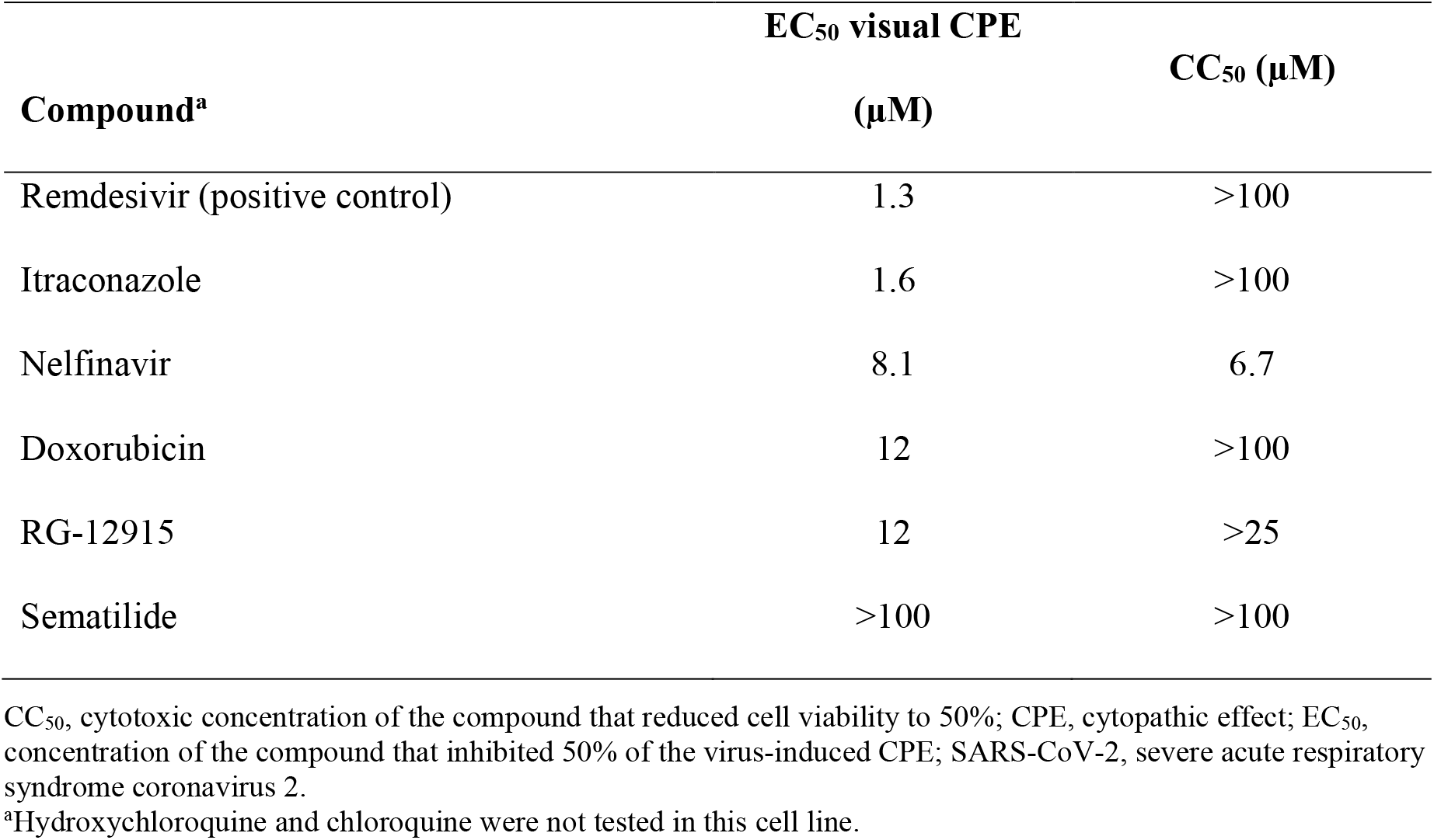
Observed antiviral activity (EC_50_) against SARS-CoV-2 and cytotoxicity (CC_50_) of the 6 compounds evaluated using the Caco-2 cell assay

| Compound<sup>a</sup> | EC<sub>50</sub> visual CPE | CC<sub>50</sub> (μM) |
| --- | --- | --- |
|  | (μM) |  |
| Remdesivir (positive control) | 1.3 | >100 |
| Itraconazole | 1.6 | >100 |
| Nelfinavir | 8.1 | 6.7 |
| Doxorubicin | 12 | >100 |
| RG-12915 | 12 | >25 |
| Sematilide | >100 | >100 |

CC<sub>50</sub>, cytotoxic concentration of the compound that reduced cell viability to 50%; CPE, cytopathic effect; EC<sub>50</sub>, concentration of the compound that inhibited 50% of the virus-induced CPE; SARS-CoV-2, severe acute respiratory syndrome coronavirus 2.
<sup>a</sup>Hydroxychloroquine and chloroquine were not tested in this cell line.

## DISCUSSION

Early in the pandemic, there was an urgent need for treatment of COVID-19, and the most rapid solution involved repurposing drugs already validated in the clinic. To accelerate drug repurposing, libraries have been screened in computational models to target viral proteins, including the receptor-binding domain of the spike protein, ACE2, viral Plpro, 3Clpro, and Mpro.^18,26^ In parallel, cell-based screening using a variety of cell lines, including Vero E6, Huh7, Calu-3, and Caco-2 cells infected with SARS-CoV-2, were developed to identify potential candidates from repurposing libraries.^9,27–30^

The drug repurposing library screen described in this study used a VeroE6-eGFP SARS-CoV-2 infection assay to identify therapeutic drug candidates. This cell line is known to be susceptible to SARS-CoV-1 and SARS-CoV-2 infection and shows a reduced eGFP signal upon cell death.^11^ The HTS screening assay developed in this study was robust (**Table I**), automated, and represents a scalable and efficient system for identifying antiviral compounds. Although some previously developed assays have used HCI,^9,27,28^ few have compared HCI and PR readout, as described in this study. In developing HTS assays, factors such as data storage, data acquisition speed, and logistics should be considered, especially when speed is of utmost importance, such as during a pandemic. In addition to the advantages of speed and storage afforded by PR, all parameters for HTS quality (RZ’, S/N), intra- and inter-plate variation were overall substantially better using HCI compared with PR readout (**Table I, Table II, Figure 2**). This made HCI the preferred readout for screening.

A library of 5676 compounds that passed phase 1 were screened. The primary hit rate was 4.0%, which is relatively high among experimental repurposing screening studies that usually report hit rates of less than 2%.^30–32^ This rather high hit rate is due to the low threshold that was set for primary hit selection and because toxicity was not taken into account for primary hit selection. In total, 52 hits were confirmed. After elimination of toxic compounds, screening artefacts (eg, vitamin B2 and orantinib cause a false positive signal due to fluorescence of the compound), and unwanted modes of action such as influence on lysosomal function or phospholipidosis,^33^ 8 compounds remained (**Table III**). Five of these compounds also exhibited antiviral activity in Caco-2 cells, a validated model for SARS-CoV research,^34^ showing that these compounds were active across different cell types.

Although hydroxychloroquine and its analogue chloroquine inhibit SARS-CoV-2 entry and replication,^35^ they failed in clinical trials.^36,37^ Of the 8 selected compounds, only remdesivir, an adenosine nucleotide prodrug that inhibits the viral RNA-dependent RNA polymerase,^18,38^ has been approved by the US Food and Drug Administration (FDA). In the current study, remdesivir consistently produced EC_50_ values of single-digit micromolar magnitude in VeroE6 cells and in Caco-2 cells; these values were comparable to those reported in previous studies (range, 0.77 µM to 7.28 µM).^9,10,25,27,29^ Remdesivir was the first drug approved by the FDA for the treatment of patients with COVID-19 requiring hospitalisation in the United States and is conditionally recommended by Infectious Diseases Society of America guidelines for use in hospitalised patients with severe COVID-19.^37^

Although examples of successful drug repurposing exist (eg, sildenafil citrate),^39^ the outcome in the case of SARS-CoV-2 and COVID-19 has been disappointing; however, there are lessons to be learned from this unprecedented effort, principally regarding assay design and contextualisation of data.^40^ Almost two years after the emergence of the first cases of COVID-19 in China, the pandemic remains ongoing, and new compounds are under investigation. AT-527, a guaniside nucleoside analogue, was one of the first new drugs to be tested in clinical trials after promising pre-clinical data^41^; however, the compound did not show a reduction from baseline in the amount of SARS-CoV-2 virus in patients with mild or moderate COVID-19 compared to placebo in the clinics.^42^ PF-07321332,^43^ an orally bioavailable Mpro inhibitor, is currently undergoing evaluation in combination with ritonavir in two trials exploring progression to severe disease in different populations (NCT04960202, NCT05011513) and a study assessing post-exposure treatment (NCT05047601). Interim analysis of the phase 2/3 Evaluation of Protease Inhibition for COVID-19 in High-Risk Patients (EPIC-HR) randomised, double-blind study of non-hospitalised adult patients with COVID-19 showed an 89% reduction in risk of COVID-19-related hospitalisation or death from any cause compared to placebo.^44^ In addition, molnupiravir (EIDD-01931), a nucleoside analogue with broad antiviral activity, was found to have good antiviral activity and favourable pharmacokinetic properties and was active *in vivo*.^45^ After finalisation of the screen, molnupiravir was tested in the VeroE6-eGFP cells and found to be active with an EC_50_ of ∼4 µM. After promising results in clinical trials (NCT04575597), molnupiravir received marketing approval in the UK, and is currently under review by the FDA and the European Medicines Agency.

## CONCLUSIONS

The findings of this study were comparable with other large screening studies that identified potential anti–SARS-CoV-2 agents and compounds, although *in vivo* efficacy, toxicity, and pharmacokinetic investigation of the selected hits in this report did not support novel application against SARS-CoV-2.^9,12,25,27,29,46–50^ Together, these studies demonstrate that drug repurposing technology can identify potential agents for rapidly spreading novel pathogens, as in the case of remdesivir, which has been used to treat patients with severe COVID-19. Although preclinical drug repurposing studies may provide a means of identifying potential agents and avoiding some of the limitations of traditional clinical development, the findings of this study suggest that results from *in silico* or *in vitro* studies should be interpreted with caution. Evaluations of compounds *in vivo* and in clinical trials are necessary to support the use of identified compounds in clinical settings.

## ACKNOWLEDGMENTS

Medical writing support for the development of this manuscript was provided by Kurt Kunz, MD, MPH, and Catherine DeBrosse, PhD, of Cello Health Communications/MedErgy, and was funded by Janssen Pharmaceutica.

## Notes

**FUNDING STATEMENT** This work has been executed as part of the Corona Accelerated R&D in Europe (CARE) project under grant agreement number 101005077 with the Innovative Medicines Initiative 2 Joint Undertaking (JU). The JU receives support from the European Union’s Horizon 2020 research and innovation programme EFPIA, Bill & Melinda Gates Foundation, Global Health Drug Discovery Institute, and the University of Dundee. The content of this publication only reflects the authors’ views and the JU is not responsible for any use that may be made of the information it contains. This project has received funding from the European Union’s Horizon 2020 research and innovation programme under grant agreement number 101003627. Part of this research was performed using the ‘Caps-It’ research infrastructure (project ZW13-02) that was financially supported by the Hercules Foundation and Rega Foundation, KU Leuven. This work has been funded in part with Federal funds from the Office of the Assistant Secretary for Preparedness and Response, Biomedical Advanced Research and Development Authority (BARDA), under OTA number HHSO100201800012C. Employees of the sponsor, Janssen Pharmaceutica NV, contributed to the study design, data analysis and interpretation, the writing of the report, and the decision to submit the manuscript for publication.

**CONFLICT OF INTEREST DISCLOSURE** The authors declare the following financial interests/personal relationships which may be considered as potential competing interests: SC received research funding from Janssen for this research. WC, DJ, SDJ, PL, and JN are employees from KU Leuven and received funding from Janssen for this research. LV, CVdE, CB, SDM, MVL, and EVD are employees of Janssen and may be stock owners of Johnson & Johnson. DB and JC have nothing to declare.

### Competing Interest Statement

The authors declare the following financial interests/personal relationships which may be considered as potential competing interests: SC received research funding from Janssen for this research. WC, DJ, SDJ, PL, and JN are employees from KU Leuven and received funding from Janssen for this research. LV, CVdE, CB, SDM, MVL, and EVD are employees of Janssen and may be stock owners of Johnson & Johnson. DB and JC have nothing to declare.

