## Supplementary material for "Development and optimisation of a high-throughput screening assay for in vitro anti–SARS-CoV-2 activity: evaluation of 5676 phase 1 passed structures": Supplemenatary Material

| Identifier | Name | GFP plate reader anti-viral screen EC50 (µM) | HCI antiviral screen EC50 (µM) | ATPlite toxicity CC50 (µM) |
|---|---|---|---|---|
| COVC-0134286843 | EVP-6124_ENCENICLINE | >100.00 | >100.00 | 20.9 |
| COVC-0135464447 | VE-822 | > 5.01 | 8.32 | 10.2 |
| COVC-0135517367 | ORANTINIB STEREOISOMER | 2 | > 50.10 | > 50.10 |
| COVC-0136083474 | HALOFANTRINE | > 20.00 | 9.12 | 15.1 |
| COVC-0136118120 | PECAZINE | > 20.00 | >100.00 | 11.8 |
| COVC-0136318398 | CANNABINOR | >100.00 | > 50.10 | 7.67 |
| COVC-0136544954 | CYCLOSPORINE | >100.00 | >100.00 | 18.6 |
| COVC-0136580428 | ALMOREXANT | 18.6 | > 25.10 | 21.4 |
| COVC-0136643402 | BEDAQUILINE | > 50.10 | > 50.10 | 13.7 |
| COVC-0138110417 | RAVIDASVIR | >100.00 | >100.00 | 12.7 |
| COVC-0139007958 | ICI-195739 | >100.00 | >100.00 | 25.7 |
| COVC-0139169083 | JNJ-18038683 | > 25.10;> 20.00 | > 25.10 | 14.6 |
| COVC-0139464749 | BP-897 | > 25.10 | > 25.10 | 30.2 |
| COVC-0139663433 | 8-AZAGUANINE | > 50.10 | > 50.10 | 23.7 |
| COVC-0140030971 | PIPENDOXIFENE | 3.63 | > 6.31 | 4.62 |
| COVC-0140330787 | TMC310911 | 6.92; 2.24 | 9.55;>100.00 | 14.30; 11.80 |
| COVC-0140549998 | SALMATEROL | 19.1 | >100.00 | 18.6 |
| COVC-0140833037 | TIOPERIDONE | 9.12 | >100.00 | 7.24 |
| COVC-0140864590 | TRIMEPRAZINE | 17.4 | > 25.10 | 21.4 |
| COVC-0140958357 | TIPIFARNIB | >100.00 | >100.00 | 13.3 |
| COVC-0141087222 | CLOMIPRAMINE | 14.5 | >100.00 | 16 |
| COVC-0141101284 | MCN-5691 | 4.27 | >100.00 | 4.84 |
| COVC-0141142856 | MEQUITAZINE | 7.76 | >100.00 | 7.76 |
| COVC-0141155358 | FLUSPIRILENE | >100.00 | >100.00 | 7 |
| COVC-0141155434 | PHENOXYBENZAMINE HYDROCHLORIDE | 20 | >100.00 | |
| COVC-0141206740 | LESOPITRON | 2.34 | >100.00 | >100.00 |
| COVC-0141212165 | NORTRIPTYLINE | 15.9 | >100.00 | 24 |
| COVC-0141261705 | EVANS-BLUE | >100.00 | >100.00 | 18 |
| COVC-0149442021 | MOCRAVIMOD | 3.31 | >100.00 | 3.06 |
| COVC-0149674334 | BREXPIPRAZOLE | >100.00 | >100.00 | 12 |
| COVC-0150976668 | | | > 20.00 | |
| COVC-0150980605 | ENECADIN | 1.74 | 21.9 | |
| COVC-0672397984 | EVP-0334 | >100.00 | >100.00 | 1.55 |
| COVC-0672954843 | DULOXETINE | >100.00; 7.24 | >100.00;> 10.00 | 17.80; 14.80 |
| COVC-0674506360 | CAMOSTAT | > 20.00 | >100.00 | |
| COVC-0675007677 | DACLATASVIR | >100.00 | >100.00 | 19.7 |
| COVC-0675868528 | METITEPINE | 7.24 | >100.00 | 7 |
| COVC-0675878035 | SABELUZOLE | 18.6 | >100.00 | 15.3 |
| COVC-0675923228 | FLUPENTHIXOL | 8.71 | >100.00 | 7.67 |
| COVC-0676534350 | BAY-B-5369 | | > 20.00 | |
| COVC-0677008167 | AZELASTINE | 13.3 | >100.00 | 17.8 |
| COVC-0677194034 | REVAPRAZAN | >100.00 | 10.5 | 11.5 |
| COVC-0677582639 | SAREDUTANT (MIXTURE?) | >100.00 | >100.00 | 23.4 |
| COVC-0677720184 | EUPROCIN | >100.00 | >100.00 | 11.4 |
| COVC-0677735491 | THIOPROPERAZINE | 13.2 | >100.00 | 6.31 |
| COVC-0677952181 | BIFEMELANE | >100.00 | 20.9 | 26.9 |
| COVC-0677955252 | METHADONE | > 25.10 | > 25.10 | 13.2 |
| COVC-0677970346 | LACIDIPINE | > 10.00 | | 7.24 |
| COVC-0677970490 | TIOSPIRONE | >100.00 | >100.00 | 24.6 |
| COVC-0678000383 | CURCUMIN | >100.00 | >100.00 | 14.8 |
| COVC-0678006086 | ABIRATERONE ACETATE | >100.00 | > 25.10 | 9.89 |
| COVC-0678009973 | SEMATILIDE | 3.89 | >100.00 | >100.00 |
| COVC-0678023366 | CLOCINIZINE | >100.00 | 20.9 | 21.9 |
| COVC-0678033956 | PROMETHAZINE | 21.9 | >100.00 | 33.1 |
| COVC-0678071379 | PIPOTHIAZINE | 13.8 | >100.00 | 7.33 |
| COVC-0678151537 | METHDILAZINE | 16.2 | >100.00 | 14.8 |
| COVC-0678170981 | INDATRALINE | 8.32 | >100.00 | 8.51 |
| COVC-0678171662 | THIOTHIXENE | >100.00 | 5.75 | 4.96 |
| COVC-0678230812 | DRAFLAZINE | >100.00 | >100.00 | 9.66 |
| COVC-0678388189 | HALETAZOLE | 3.89 | >100.00;> 10.00 | 3.72 |
| COVC-0678849574 | SEPROXETINE | 7.59 | >100.00 | 10.6 |

| Identifier | Name | GFP plate reader anti-viral screen EC50 (µM) | HCI antiviral screen EC50 (µM) | ATPlite toxicity CC50 (µM) |
|---|---|---|---|---|
| COVC-0678867755 | BENIDIPINE | >100.00 | >100.00 | 14.8 |
| COVC-0678879901 | ITRACONAZOLE | >100.00 | | >100.00 |
| COVC-0686249369 | TAK-652 | | > 20.00 | |
| COVC-0687833464 | | | > 20.00 | |
| COVC-0687841008 | | | > 20.00 | |
| COVC-0687843163 | | | > 20.00 | |
| COVC-0687847569 | | | > 20.00 | |
| COVC-0687852474 | BGP-15 | 0.525 | >100.00 | |
| COVC-0687853800 | EIDD-2801 | 14.1 | 4.43 | |
| COVC-0687854362 | ANAMORELIN HYDROCHLORIDE | | > 20.00 | |
| COVC-1207972476 | DR1289116 | >100.00 | > 50.10 | 16 |
| COVC-1207984410 | ANDROGRAPHOLIDE | >100.00 | >100.00 | 21.6 |
| COVC-1208049825 | GS-938 | > 20.00 | > 50.10 | |
| COVC-1210360354 | METHOTRIMEPRAZINE_LEVOMEPROMAZINE | 11.8 | > 50.10 | 11.6 |
| COVC-1210360383 | MONENSIN | > 20.00 | 1.12 | |
| COVC-1210363295 | PRENYLAMINE | 8.71 | >100.00 | 8.41 |
| COVC-1210363846 | SULCOTIDIL | >100.00 | >100.00 | 3.27 |
| COVC-1211756174 | DARIFENACIN | > 25.10 | >100.00 | 56.2 |
| COVC-1212730902 | NEBIVOLOL | 8.91 | >100.00 | 7 |
| COVC-1212741697 | R-167154 ISOMER | >100.00 | >100.00 | 19.5 |
| COVC-1212748902 | NEBIVOLOL | > 10.00 | >100.00 | 8.71 |
| COVC-1213406057 | PIPERLONGUMINE | > 50.10 | > 50.10 | 9.55 |
| COVC-1214272359 | CLEMASTINE | > 50.10 | 20.4 | 18.4 |
| COVC-1214358711 | BROFAROMINE | 14.8 | >100.00 | 19.1 |
| COVC-1214407107 | QUININE_QUINIDINE | 15.1 | > 25.10 | > 25.10 |
| COVC-1214563522 | BITOLTEROL | 8.13 | >100.00 | 5.69 |
| COVC-1214565530 | METIXENE | 9.55 | >100.00 | 10.1 |
| COVC-1214607586 | PROPIPOCAINE | >100.00 | >100.00 | 31.6 |
| COVC-1214614542 | CARFENAZINE | >100.00 | >100.00 | 6.31 |
| COVC-1214621709 | FENOBAM | > 20.00 | > 25.10 | |
| COVC-1214767323 | ZOTEPINE | >100.00 | >100.00 | 12.3 |
| COVC-1214826443 | GIRISOPAM | >100.00 | >100.00 | >100.00 |
| COVC-1214829248 | HALOFANTRINE | 9.55 | 9.12 | 11.5 |
| COVC-1214857454 | FLUPHENAZINE | >100.00 | >100.00 | 7.76 |
| COVC-1214857892 | OXATOMIDE | >100.00 | >100.00 | 12.6 |
| COVC-1214895840 | SPIPERONE | 20.4 | 22.4 | 81.3 |
| COVC-1214897179 | PENFLURIDOL | >100.00 | >100.00 | 3.59 |
| COVC-1214903948 | INDOMETACIN | >100.00 | >100.00;> 50.10 | >100.00 |
| COVC-1215023048 | METHYPRYLON | 18.4 | > 25.10 | > 25.10 |
| COVC-1215027317 | LEVOMEPROMAZINE | >100.00 | >100.00 | 22.9 |
| COVC-1215738110 | CLOMIFENE | > 20.00 | > 20.00 | |
| COVC-1222873116 | HYDROXYCHLOROQUINE STEREOISOMER(S) | 12.7 | 18.8 | > 50.10 |
| COVC-1223085504 | CCS-1477 (S*) | >100.00 | >100.00 | 3.16 |
| COVC-1224702964 | | | >100.00 | |
| COVC-1746032955 | REMDESIVIR | 4.64 | 3.12 | 95.5 |
| COVC-1746696265 | ATRACURIUM BESYLATE | | > 20.00 | |
| COVC-1746967931 | TOREFORANT | > 25.10 | > 50.10 | 22.4 |
| COVC-1747138106 | BERBAMINE | 8.51 | > 6.31 | 10.2 |
| COVC-1747138440 | SALINOMYCIN | > 20.00 | 1.29 | |
| COVC-1747138479 | BEBEERINE | > 10.00 | 5.37 | 11.5 |
| COVC-1747162848 | CASOPITANT | > 25.10 | > 25.10 | 17.8 |
| COVC-1747188285 | BI-2536 | > 50.10 | > 50.10 | 0.603 |
| COVC-1747231597 | DEPTROPINE | 7.41 | > 50.10 | 13.5 |
| COVC-1747292777 | LYNESTRENOL | > 50.10 | > 50.10 | 10.5 |
| COVC-1747397988 | HT-61 | > 20.00 | 4.9 | |
| COVC-1748269057 | NERATINIB | | > 10.00 | |
| COVC-1751145778 | DITHRANOL | >100.00 | >100.00 | >100.00 |
| COVC-1751409823 | SYROSINGOPINE | >100.00 | >100.00 | 8.91 |
| COVC-1751415324 | OXAPROZIN | 11.8 | > 50.10 | > 50.10 |
| COVC-1751439919 | PERAZINE | 14 | >100.00 | 14.6 |
| COVC-1751458750 | NOXIPTILINE | 14.1 | >100.00 | 77.6 |

| Identifier | Name | GFP plate reader anti-viral screen EC50 (µM) | HCI antiviral screen EC50 (µM) | ATPlite toxicity CC50 (µM) |
|---|---|---|---|---|
| COVC-1751645509 | CARPIPRAMINE | 8.91 | >100.00 | 8.61 |
| COVC-1751700625 | CLOPIPAZAN | > 20.00 | >100.00 | 15 |
| COVC-1751748815 | METHOXYTACRINE | 11.5 | > 25.10 | 47.9 |
| COVC-1751763386 | PROADIFEN | 5.13 | >100.00 | 24 |
| COVC-1751764998 | ASTEMIZOLE | >100.00 | >100.00 | 7.5 |
| COVC-1751771027 | PROFLAVINE | 15 | > 25.10 | 6.61 |
| COVC-1751892074 | BBR-3464 | | > 20.00 | |
| COVC-1751893009 | DIPERODON | 15.1 | > 25.10 | > 25.10 |
| COVC-1752607069 | SAPERCONAZOLE | >100.00;> 20.00 | >100.00 | >100.00 |
| COVC-1752620251 | 3 ALLYLFENTANYL | 4.37 | 4.9 | |
| COVC-1760096432 | GSK-921 | > 50.10 | > 50.10 | 2.63 |
| COVC-1760099291 | AJULEMIC ACID | >100.00 | >100.00 | 21.4 |
| COVC-1760287043 | SULFATINIB | > 10.00 | > 50.10 | 4.9 |
| COVC-1761575739 | | | > 20.00 | |
| COVC-1761588262 | | | > 20.00 | |
| COVC-1761590281 | | | > 20.00 | |
| COVC-1761595595 | V-GLYCOPEPTIDE | | > 20.00 | |
| COVC-1761604319 | ADOMEGLIVANT | > 10.00 | > 6.31 | 10.4 |
| COVC-2282947189 | VALRUBICIN | | > 20.00 | |
| COVC-2283008312 | NAVITOCLAX | >100.00 | >100.00 | 4.73 |
| COVC-2283590314 | OSIMERTINIB | >100.00 | >100.00 | 3.09 |
| COVC-2283593123 | GSK1070916 | >100.00 | >100.00 | 9.33 |
| COVC-2285168592 | VINORELBINE | | > 20.00 | |
| COVC-2285169276 | C-172 | 19.5 | 14.5 | 28.2 |
| COVC-2286471467 | LOFENTANIL | 20.9 | > 25.10 | |
| COVC-2286484880 | METHYLDOPA | 17.8 | >100.00 | |
| COVC-2286491085 | NEBIVOLOL | >100.00 | >100.00 | 10.2 |
| COVC-2286528427 | CDP-840 (B) | >100.00 | >100.00 | 23.4 |
| COVC-2286535978 | PENBUTOLOL | 15.1 | 20 | 29.5 |
| COVC-2287142567 | NORCYCLOBENZAPRINE | > 50.10 | > 50.10 | 23.4 |
| COVC-2287147580 | CEFOPERAZONE | > 20.00 | > 20.00 | |
| COVC-2287807964 | SAQUINAVIR | >100.00 | >100.00 | 18.2 |
| COVC-2288269965 | PIPERACETAZINE | 16.2 | > 12.60 | 12.2 |
| COVC-2288318596 | METHOPROMAZINE | > 20.00 | >100.00 | 14.1 |
| COVC-2288327793 | DIAMOCAINE | 9.55 | 22.4 | 46.8 |
| COVC-2288339516 | OXETORONE | 18.2 | >100.00 | 12.2 |
| COVC-2288570388 | DOSULEPIN | 16.6 | > 25.10 | 20 |
| COVC-2288571827 | BUCINDOLOL | 17 | >100.00 | 30.9 |
| COVC-2288600616 | DRONEDARONE | 3.02 | >100.00 | 3.85 |
| COVC-2288609209 | LOPERAMIDE | 11.8 | 8.91 | 22 |
| COVC-2288621041 | PRENYLAMINE | 8.51 | >100.00 | 10.7 |
| COVC-2288636389 | ZIMELIDINE | > 20.00 | | > 10.00 |
| COVC-2288688211 | MILENPERONE | >100.00 | >100.00 | 32.4 |
| COVC-2288695035 | CARVEDILOL | 15.5 | 10 | 17 |
| COVC-2288727795 | NAFOXIDINE | >100.00 | >100.00 | 3.85 |
| COVC-2288809083 | RHODOQUINE | 24.6 | >100.00 | >100.00 |
| COVC-2288955269 | LIFIBRATE | >100.00 | >100.00 | 38 |
| COVC-2289454566 | BEDAQUILINE | >100.00 | 15.5 | 26.9 |
| COVC-2289479927 | ZUCLOPENTHIXOL_CLOPENTHIXOL | 7.24 | >100.00 | 6.68 |
| COVC-2289732646 | PIMETHIXENE | >100.00 | >100.00 | 25.7 |
| COVC-2298439635 | | | > 20.00;> 20.00 | |
| COVC-2298460306 | | | > 20.00 | |
| COVC-2298477237 | O-304 | | > 20.00 | |
| COVC-2819871799 | IVACAFTOR | 4.57 | > 25.10 | 4.62 |
| COVC-2820438510 | LOTEPREDNOL ETABONATE | | > 20.00 | |
| COVC-2820464103 | VOLASERTIB | > 50.10 | > 50.10 | 0.234 |
| COVC-2820786116 | TETRANDRINE_HANFANGCHIN A | 3.98 | > 50.10 | 5.17 |
| COVC-2820802843 | CRIZOTINIB | 6.76 | 4.47 | 4.42 |
| COVC-2821037923 | FLUORESCEIN | | >0.3090 | > 20.00 |
| COVC-2821711673 | HYDROQUINIDINE | 15.5 | > 25.10 | > 25.10 |
| COVC-2822096521 | RENYTOLINE | 3.47 | >100.00 | 2.92 |

| Identifier | Name | GFP plate reader anti-viral screen EC50 (µM) | HCI antiviral screen EC50 (µM) | ATPlite toxicity CC50 (µM) |
|---|---|---|---|---|
| COVC-2823343639 | NEBIVOLOL | >100.00 | >100.00 | 7.85 |
| COVC-2823361537 | NEBIVOLOL | >100.00 | >100.00 | 7.76 |
| COVC-2824600869 | ARIPIPRAZOLE | >100.00 | >100.00 | 11.4 |
| COVC-2825210622 | BUCLIZINE | >100.00 | 20.4 | 81.3 |
| COVC-2825221500 | SERCLOREMINE | 17.2 | >100.00 | 34.7 |
| COVC-2825327487 | PAROXETINE | 15.1 | >100.00 | 16.6 |
| COVC-2825450452 | CLOSPIPRAMINE | 7.41 | >100.00 | 8.13 |
| COVC-2825472328 | LIDOFLAZINE | 9.55 | >100.00 | 8.41 |
| COVC-2825487070 | ARBIDOL | >100.00 | >100.00 | |
| COVC-2825507141 | ACEPROMAZINE | >100.00 | >100.00 | 22.4 |
| COVC-2825508294 | MASOPROCOL | >100.00 | >100.00 | 37.2 |
| COVC-2825514435 | CHLOROQUINE | 5.86 | 9.46 | 38.7 |
| COVC-2825515130 | QUINESTROL | 8.71 | >100.00 | 7.59 |
| COVC-2825616197 | CLOBENZTROPINE | 9.77 | >100.00 | 8.13 |
| COVC-2825616302 | DIHYDREXIDINE | 7.94 | >100.00 | 18.6 |
| COVC-2825622218 | OXETHAZAINE | 18.6 | >100.00 | 26.3 |
| COVC-2825640648 | HOMOCHLORCYCLIZINE | > 20.00 | > 25.10 | 21.1 |
| COVC-2825645391 | TERFENADINE | >100.00 | >100.00 | 5.13 |
| COVC-2826330726 | ZAMIFENACIN | >100.00 | 87.1 | >100.00 |
| COVC-2826351877 | SABELUZOLE | >100.00 | >100.00 | 15.5 |
| COVC-2833485378 | HYDROXYCHLOROQUINE (R*) | 12.7 | > 50.10 | > 50.10 |
| COVC-2833739552 | QCC-374 | 17.4 | 16.6 | 50.1 |
| COVC-2835343357 | LEVOMEFOLATE | | > 20.00 | |
| COVC-3356547646 | HDAC-42_AR-42 | > 20.00 | | 0.851 |
| COVC-3356682871 | TEPOTINIB_EMD-1214063 | >100.00 | >100.00 | 5.75 |
| COVC-3356809212 | CYTARABINE | > 50.10 | > 20.00 | |
| COVC-3357305447 | TPEN | > 3.09 | >100.00 | 1.93 |
| COVC-3357309398 | CEFUROXIME AXETIL | | > 20.00 | |
| COVC-3357775383 | SUDOTERB | >100.00 | >100.00 | 14.3 |
| COVC-3357844334 | TRIBENOSIDE | > 50.10 | > 50.10 | 17.4 |
| COVC-3358854404 | PONATINIB_ICLUSIG | > 20.00;>100.00 | >100.00 | 0.9890;< 1.55 |
| COVC-3359139833 | ERDAFITINIB | > 10.00;>100.00 | > 6.31 | 5.25; 3.63 |
| COVC-3359217199 | MK-0893 | >100.00 | >100.00 | 18.8 |
| COVC-3360221789 | EPIRUBICIN | | >100.00 | |
| COVC-3360223611 | CINNARIZINE | 18.2 | >100.00 | 14.5 |
| COVC-3360248091 | DOXORUBICIN | 8.04 | 33.1 | 64.6 |
| COVC-3360252970 | ALVOCIDIB | | > 10.00 | |
| COVC-3360889795 | CONESSINE | 12 | > 25.10 | 34.7 |
| COVC-3361592863 | TMC-353121 | > 20.00 | > 20.00;> 5.01 | 9.33 |
| COVC-3361708049 | AMG-487-(+/-) | > 20.00 | > 20.00 | |
| COVC-3361719627 | TANDUTINIB | >100.00 | >100.00 | 11.5 |
| COVC-3362013046 | PERICIAZINE | > 20.00 | >100.00 | 12.7 |
| COVC-3362049948 | METOFENAZATE | 18.6 | >100.00 | >100.00 |
| COVC-3362050611 | ACETOPHENAZINE | >100.00 | >100.00 | 14.6 |
| COVC-3362302644 | FLUPIRTINE | >100.00 | >100.00 | >100.00 |
| COVC-3362312915 | PROTRIPTYLINE | >100.00 | >100.00 | 26.3 |
| COVC-3362312957 | THIORIDAZINE | 6.76 | >100.00 | 6.38 |
| COVC-3362314340 | BENZATROPINE | > 25.10 | >100.00 | 46.8 |
| COVC-3362314676 | RAUNORMINE_DESERPIDINE | > 20.00 | > 25.10 | 11.5 |
| COVC-3362317361 | QUINESTROL | >100.00 | >100.00 | 8.22 |
| COVC-3362343100 | TERFLAVOXATE | >100.00 | >100.00 | 63.1 |
| COVC-3362347376 | TRIFLUOPERAZINE | 6.46 | >100.00 | 7.94 |
| COVC-3362350497 | CHLORBENZOXAMINE | >100.00 | > 25.10 | 15.3 |
| COVC-3362350605 | LASALOCID | >0.6310 | 1.07 | 5.37 |
| COVC-3362375297 | NORGESTIMATE | >100.00 | >100.00 | 20.9 |
| COVC-3362380349 | CINCHOCAINE | 29.1 | 29.5 | 39.6 |
| COVC-3362381936 | PROMAZINE | 16.2 | >100.00 | 25.7 |
| COVC-3362486392 | DOTARIZINE | > 20.00 | 10 | 17.4 |
| COVC-3363187991 | COLCHICINE | >100.00 | >100.00 | |
| COVC-3370896893 | R-428_BEMCENTINIB | 1.78 | > 50.10 | 1.55 |
| COVC-3372196066 | ROVAZOLAC | | > 20.00 | |

| Identifier | Name | GFP plate reader anti-viral screen EC50 (µM) | HCI antiviral screen EC50 (µM) | ATPlite toxicity CC50 (µM) |
|---|---|---|---|---|
| COVC-3372202143 | | | > 20.00 | |
| COVC-3372208332 | L-1777 | | > 20.00 | |
| COVC-3372208930 | MAVORIXAFOR | | > 20.00 | |
| COVC-3893557264 | BMS833923 | 2.95 | >100.00 | 3.09 |
| COVC-3893559927 | RIBOCICLIB | >100.00 | >100.00 | 38.9 |
| COVC-3894275940 | TECALCET | 16.6 | >100.00 | 28.2 |
| COVC-3894718404 | DOSULEPIN | 14.3 | > 25.10 | 17 |
| COVC-3894856214 | GIVINOSTAT | 10.5 | >100.00 | 0.767 |
| COVC-3895781309 | ALOXISTATIN | 34.3 | 85.1 | >100.00 |
| COVC-3897093565 | NEBIVOLOL | >100.00 | >100.00 | 7.16 |
| COVC-3897104103 | SABELUZOLE | 18.6 | >100.00 | 19.5 |
| COVC-3897760602 | VITAMINE B2_RIBOFLAVIN | 11.2 | > 50.10 | > 50.10 |
| COVC-3898089676 | DARUNAVIR | > 20.00 | > 20.00 | |
| COVC-3898253804 | LOPINAVIR | 22.9 | 15.7 | 33.7 |
| COVC-3898420702 | NELFINAVIR | 5.22 | 3.50; 4.10 | 11.2 |
| COVC-3898606637 | JNJ-5207852 | > 20.00 | > 20.00 | |
| COVC-3898674512 | IMATINIB_GLEEVEC_ | 11 | 21.9 | 11.9 |
| COVC-3898855212 | CHLORPROTHIXENE | 7.08 | >100.00 | 13.8 |
| COVC-3898932127 | BUTOPROZINE | >100.00 | >100.00 | 6.61 |
| COVC-3898935732 | OXAPROTILINE | >100.00 | >100.00 | 28.2 |
| COVC-3898956268 | CLOXACEPRIDE | 3.63 | >100.00 | 2.85 |
| COVC-3899173551 | FLUTROLINE | 4.68 | >100.00 | 6.92 |
| COVC-3899185596 | RESCINNAMINE | > 50.10 | >100.00 | 5.5 |
| COVC-3899220261 | FENRETINIDE | >100.00 | 2.95 | 5.01 |
| COVC-3899221661 | RESERPINE | > 25.10 | > 50.10 | 11 |
| COVC-3899227081 | ALGESTONE ACETOPHENIDE | >100.00 | > 12.60 | 15.5 |
| COVC-3899229498 | RG-12915 | 7.94 | > 25.10 | 21.6 |
| COVC-3899304998 | FEZOLAMINE | 14.5 | >100.00 | 39.8 |
| COVC-3899357267 | DILAZEP | >100.00 | >100.00 | 28.2 |
| COVC-3899382603 | HYDROXYCHLOROQUINE | 11.6 | > 10.00; 18.00 | 47.9 |
| COVC-3899396989 | CYCLOSPORINE | 6.61 | > 50.10 | 26.3 |
| COVC-3899449667 | SEGANSERIN | 7.59 | >100.00 | 10.2 |
| COVC-3900064416 | ASENAPINE | > 25.10 | >100.00 | 5.89 |
| COVC-3900085863 | POSACONAZOLE | >100.00 | >100.00 | >100.00 |
| COVC-3900086865 | FLUOTRACEN | >100.00 | 10.5 | 12 |
| COVC-3900092763 | NABILONE | 8.91 | > 25.10 | 11.1 |
| COVC-3900101626 | TERCONAZOLE | 16.6 | > 12.60 | 10 |
| COVC-3900102081 | RITONAVIR | > 50.10 | | 37.2 |
| COVC-3900128072 | R-167154 | >100.00 | > 50.10 | 22.4 |
| COVC-3900346251 | ORANTINIB | 1.78 | > 25.10 | > 25.10 |
| COVC-3907440078 | CCS-1477 (R*) | >100.00 | >100.00 | 9.12 |
| COVC-3907758471 | ANLOTINIB | >100.00 | >100.00 | 5.25 |
| COVC-3909057522 | | | > 20.00 | |
| COVC-3909068638 | | | > 20.00 | |
